# From heterogeneous datasets to predictive models of embryonic development

**DOI:** 10.1101/2021.01.31.429006

**Authors:** Sayantan Dutta, Aleena L. Patel, Shannon E. Keenan, Stanislav Y. Shvartsman

## Abstract

Modern studies of embryogenesis are increasingly quantitative, powered by rapid advances in imaging, sequencing, and genome manipulation technologies. Deriving mechanistic insights from the complex datasets generated by these new tools requires systematic approaches for data-driven analysis of the underlying developmental processes. Here we use data from our work on signal-dependent gene repression in the *Drosophila* embryo to illustrate how computational models can compactly summarize quantitative results of live imaging, chromatin immunoprecipitation, and optogenetic perturbation experiments. The presented computational approach is ideally suited for integrating rapidly accumulating quantitative data and for guiding future studies of embryogenesis.

## Main Text

Modern studies of embryonic development started in the 1980s, when mutagenesis screens in the fruit fly revealed associations between individual genes and external structures of the larva, which hatches approximately one day after egg fertilization [1]. Most genes identified by genetic approaches encode evolutionarily conserved transcription factors and signaling enzymes. While early studies of these genes relied mainly on qualitative approaches, the ongoing research is increasingly quantitative, harnessing advances in imaging, sequencing, and genome editing [2–5]. Extracting mechanistic insights from the heterogeneous datasets generated by these new techniques, each of which reveals a different aspect of development, requires systematic strategies for data integration and analysis [6–11]. Computational models can serve as platforms for quantitative evaluation of candidate mechanisms of embryonic development and should begin to be viewed as compact repositories of data [12]. Here we illustrate this point, using datasets from genetic, biochemical, and imaging studies of the signaling enzyme Extracellular signal Regulated Kinase (ERK) and its substrate, a transcription factor Capicua (Cic), both of which are critical for normal embryogenesis and are deregulated in human diseases.

Cic is a gene repressor that controls its targets by binding to specific sequences in their regulatory DNA. Repression is relieved when Cic is phosphorylated by ERK, an enzyme activated by signaling from cell surface receptors [13–15]. ERK counteracts repression by Cic by controlling its DNA binding as well as its sub-cellular localization. The relative contributions of each mechanism to transcription remain unclear, mainly because of the differences among the experimental systems and lack of a quantitative framework for data analysis. The fruit fly embryo, a versatile developmental system where Cic was originally identified in one of the mutagenesis screens, offers unique opportunities for dissecting how distinct regulatory steps affect gene transcription. The embryo uses ERK to pattern the terminal structures of the larva and can be used to study gene regulation at multiple levels of biological organization, from molecular interactions to tissue morphogenesis [16].

We have established quantitative approaches for live imaging of Cic and its transcriptional effects in the embryo (Fig. 1c, 1e, and Supplementary Figure 1). These real-time studies reveal dynamics of molecular and cellular processes at single-cell resolution and can be coupled with biochemical assays that quantify concentration and genome-wide DNA binding patterns of Cic [17] (Fig. 1d). The power of imaging and biochemical assays is further extended by combining them with optogenetic perturbations of the signaling cascade that culminates in ERK activation [17–19]. Here we demonstrate that datasets emerging from such integrative studies (Fig. 1a) are already sufficient for constraining the parameters in a computational model that accounts for the key dynamic processes in the early embryo. The resulting model provides insights into the relative time scales and importance of the processes through which ERK relieves gene repression by Cic.

**Figure 1:**
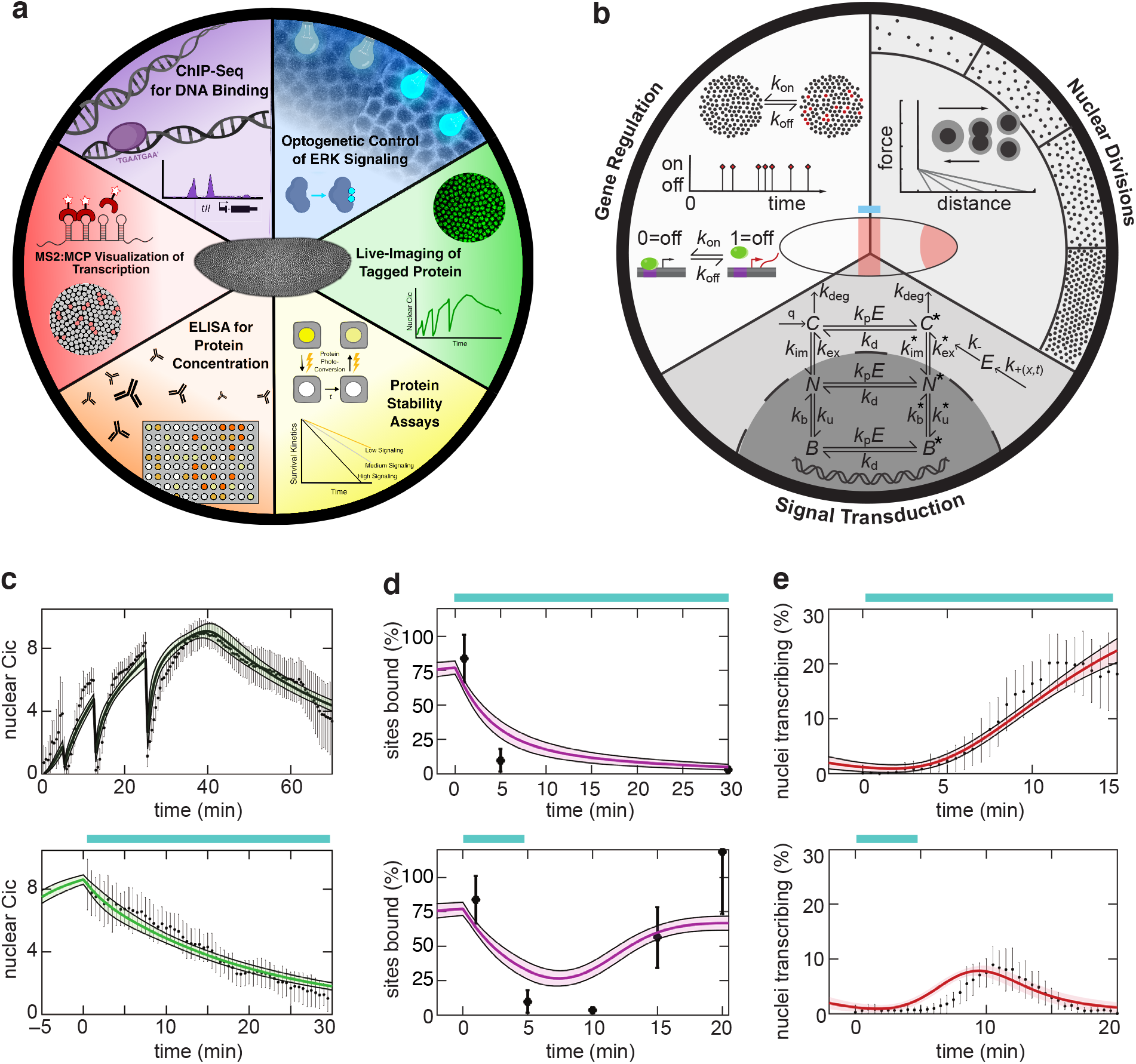
(a) A schematic of the experiments that led to the datasets used to construct the model: Ectopic activation of ERK signaling by locally activating SOS; Live imaging of fluorescently tagged Cic (green); estimation of Cic lifetime, using Cic tagged with a photoswitchable protein; ELISA measurement of molar concentration of Cic estimates the number of Cic molecules inside the nucleus; visualization of transcription via MS2-MCP reporters in the nuclei (red), respectively; measurement of the binding affinity of Cic to DNA by ChIP-Seq/ChIP-qPCR. (b) A schematic of the three tiers the model: A pairwise force-field model of nuclear divisions; a kinetic model describing the subcellular localization and biochemical state of Cic; a two-state model describes how a Cic target genes switches between the active (red) and inactive black) states. (c) Dynamics of nuclear Cic in the middle of a wild type embryo in cycle 11-14 (top) and in an embryo stimulated with optogenetic ERK signals in cycle 14 (bottom) for the simulation (green) compared to real data from imaging fluorescent protein (black) (*N* = 3 samples for both the experiments). (d) The dynamics of the fraction of Cic bound to DNA for in response to step (top) and pulsed (bottom) optogenetic ERK signals. Results from the simulation (purple) are compared to chromatin immunoprecipitation data (black) (e) Expression of the transcriptional reporter of a Cic target gene (tailless) in the middle of the embryo induced by a step (top) and pulsed (bottom) optogenetic ERK signals. Fraction of transcriptionally active nuclei in the model (red) and real embryos (black). The cyan lines in the top plots refer to start of the optogenetic stimulation. The solid colored lines show the mean simulation output from the 1000 parameter sets obtained from MCMC simulations; shaded regions represent one standard deviation from the mean. In Fig. 1c-e the black dots denote experimental mean and the errorbars denote the standard deviation across all experiments.

Our computational model is based on a three-tiered biophysical description of the following processes: nuclear divisions that establish a random nuclear packing under the common plasma membrane of the early embryo, signal-dependent nucleocytoplasmic shuttling, and transcription (Fig. 1b). Cic is synthesized in the cytoplasm and shuttles in and out of the nuclei in a tissue where the nucleocytoplasmic partitioning evolves with mitotic divisions. The rate of shuttling depends on Cic phosphorylation, which is in turn controlled by ERK. When in the cytoplasm, Cic is degraded with first order kinetics; in the nucleus, Cic reversibly binds to DNA, with phosphorylation-dependent rate constants. The Cic target gene tailless (*tll*) switches between transcriptionally active and inactive states. The integrated model has both stochastic and deterministic components: dissipative particle dynamics that produce nuclear packings (tier 1) [20], mean field kinetics with individual parameters describing nucleocytoplasmic transport, DNA binding and phosphorylation (tier 2), and a two-state Markov model for expression of the Cic target genes (tier 3) [21–23] (Methods).

Since nuclear divisions are unaffected by nucleocytoplasmic shuttling and transcription, parameters for the first tier of the model can be constrained independently of the other two, by matching the experimental and predicted statistics for internuclear distances [20]. The remaining part of the model has 12 free parameters (Supplementary Table 1). We used a stochastic optimization approach to obtain an ensemble of parameter vectors that minimize the mismatch between model predictions and datasets from measurements of Cic dynamics and its effects on transcription ([24], Methods). Data from live imaging of nuclear Cic (Supplementary Figure 1) and time resolved chromatin immunoprecipitation measurements of Cic binding to the regulatory DNA [17] constrained the parameters related to tier 2 of the model (i.e. phoshphorylation, nucleocytoplasmic shuttling and DNA binding). Parameters for the transcription part of the model were constrained by live imaging of nascent mRNA production [17] controlled by the regulatory region of *tll*, visualized with single nucleus resolution via the MS2-MCP system. All of these measurements were done in both unperturbed embryos and embryos stimulated with optogenetically induced steps and pulses of ERK activation.

Our computational model is quantitatively consistent with the most salient features of Cic regulation and function, revealed by imaging and biochemical assays (Fig. 1c-e, Supplementary Figure 2). The individual parameters of the model, most notably those related to the time scales of Cic nucleocytoplasmic shuttling and Cic-DNA binding, as well as parameters for transcription kinetics are either well constrained or have well defined upper bounds (Fig. 2a-d, Supplementary Figure 3). Furthermore, the estimated parameters quantitatively show that the Cic-DNA binding and unbinding are faster than nucleocytoplasmic shuttling of Cic – the two processes through which ERK signals control gene regulation (Supplementary Figure 4). Consistent with these findings, model predictions over the parameter ensemble show that while changes in the nuclear localization of Cic are not essential for relieving its repressor function, they provide quantitative control of the transcriptional output on a longer timescale (Fig. 2e-g). These results demonstrate how our computational model can generate quantitative predictions for the outcomes of perturbations that can not be realized experimentally, such as selective disruption of Cic nuclear import or export.

**Figure 2:**
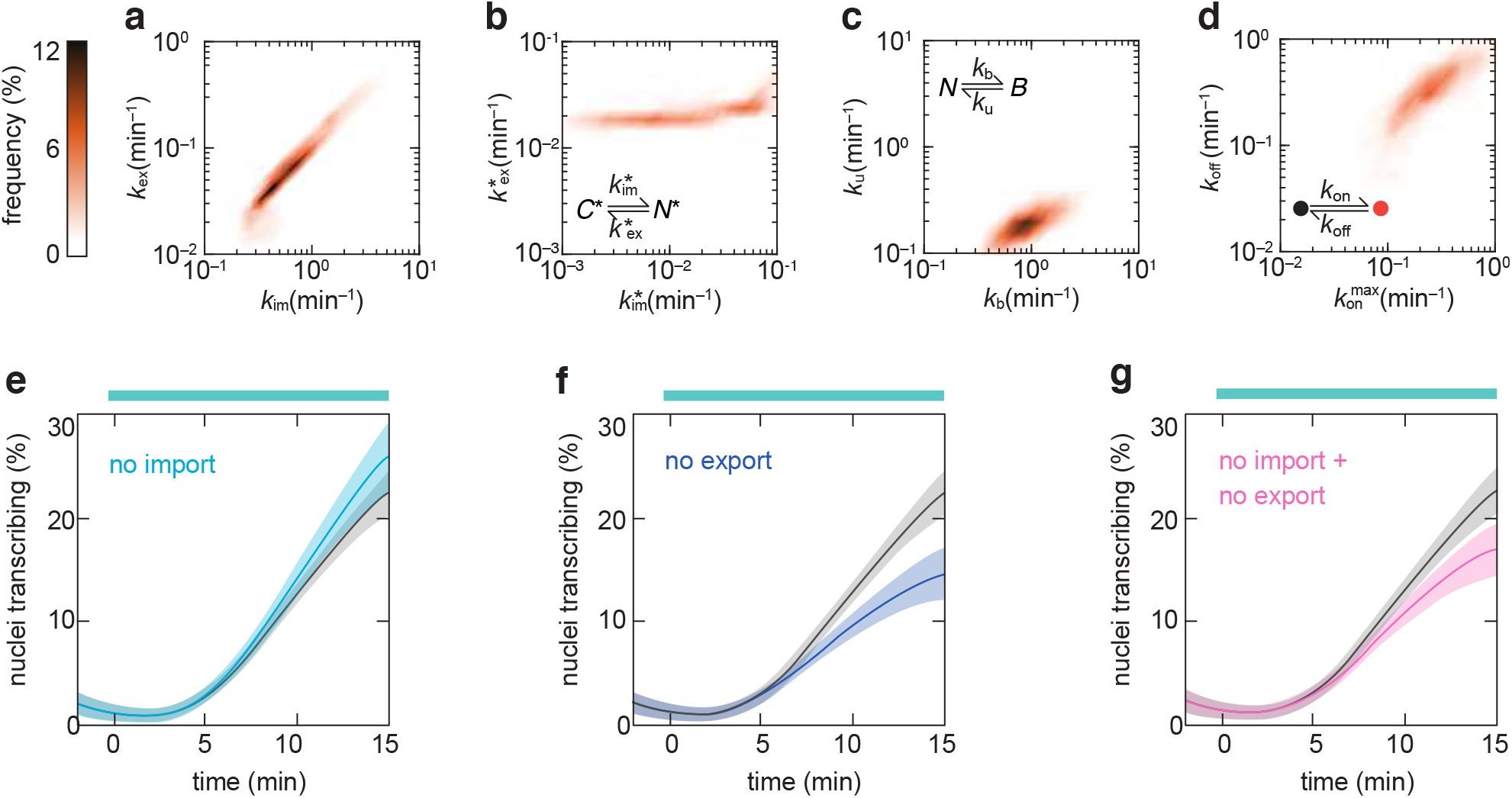
Heatmaps of the rate constants of import and export of unphosphorylated (a) and phosphorylated (b) Capicua, the rate constants of Cic-DNA binding and unbinding (c), and the rates of the activation and deactivation of the Cic target gene *tailless* (*tll*) (d). The axes are on a logarithmic scale. The color represents the probability of the respective pair of data points to be in a particular bin on the 2D plot. The insets show the corresponding processes. Transcriptional responses in the middle of the embryo in the full model (black lines) (e-g) compared to the models with inhibited import (light blue lines) (e), inhibited export (dark blue lines) (f), and inhibited import and export (pink lines) (g), after the ERK signal was turned on (cyan bar) in cycle 14. In each plot, the dark line represents the mean and the light shaded region around it represents a region of one standard deviation across the mean. In the absence of all nucleocytoplasmic transport, transcription is reduced slightly (g), whereas in the absence of only export it is reduced significantly (f) and in the absence of only import it is increased slightly (e).

In conclusion, the rapid convergence of time-resolved data, advanced genetic perturbations, and biophysical modeling have reached the point when models can make credible quantitative predictions regarding multiple aspects of embryogenesis, especially those that are too costly and difficult to explore experimentally. The presented framework is ideally suited for rigorous evaluation of the mechanistic consistency of heterogeneous datasets. For example, the model trained on data obtained for a known Cic target is also consistent with a fully synthetic reporter (Supplementary Figure 5). Our strategy can be used for evaluating how each individual data point within a dataset constrains the model parameters (Supplementary Figure 6) and is readily applicable to models that account for transcriptional feedbacks [25]. Finally, while our work was based on data from a single laboratory, future efforts should focus on integrating data from independent laboratories, similar to what has been successfully accomplished by the “data collaboration” framework in combustion research [12]. If successful, such a framework should lead to more efficient, cost-effective and, ultimately, more insightful studies of embryonic development.

## Methods

### Cic-sfGFP live imaging

To track endogenous nuclear Cic intensity in the middle of the embryo, flies with Cic Crispr-tagged with sfGFP (previously described in [17]) were crossed to flies containing a nuclear marker, histone-RFP. Embryos from this cross have maternally deposited Cic-sfGFP and his-RFP. Collected embryos were manually dechorionated and mounted onto a slide. The slide was placed on a live imaging chamber with halocarbon oil. All imaging was done on a Leica SP5 point scanning confocal microscope using the 63X oil objective. A central region of the embryo was imaged encompassing about 500 nuclei using 20 % and 10 % laser powers for the 488 nm and 561 nm lasers respectively. A Z-stack of 12 images (step size of 1 micron) was captured every 30 seconds from nuclear cycle 12 through to gastrulation. From the red channel capturing the nuclear marker, we segmented the images using the imbinarize function of the MATLAB and found the nuclear Cic levels by averaging the intensity from the green channel capturing the sf-GFP.

To track how quickly Cic leaves the nucleus in response to a light stimulus (in other words, ERK activation), we imaged embryos expressing Cic-sfGFP and OptoSOS [26]. OptoSOS expresses SOS tagged with mCherry, indicating that a red histone marker could not be used in this case. SOS is an enzyme that catalyzes the modification of a membrane-tethered pathway component, Ras, that favors its active conformation. Once active, Ras itself catalyzes the activation of a phosphorylation cascade resulting in activated ERK, which de-represses Cic. Blue light activates Opto-SOS, so simply imaging Cic-sfGFP with the 488 laser activates ERK. Within the first 5 minutes of nuclear cycle 14 we imaged Cic-sfGFP (and subsequently activated the optogeneic construct) using 20 % 488 nm laser. A Z-stack of 12 images (step size of 1 micron) was captured every 30 seconds until gastrulation. From the OptoSOS-mcherry channel, we segmented the images using the imbinarize function of the MATLAB and identify the pixels corresponding to the membranes. We assumed all other pixels correspond to nuclei (in nuclear cycle 14 nuclei are tightly packed in the cell) and found the the nuclear Cic levels by averaging the intensity from the channel capturing the sf-GFP.

### Model Description

In our model (Fig. 1b, Supplementary Table 1), Capicua (Cic) molecules have three different localizations– cytoplasmic (*C, C*^*∗*^), nuclear but unbound to the DNA (*N, N* ^*∗*^) and nuclear and bound to the DNA (*B, B*^*∗*^). The molecule in each localization can be in two states– unphosphorylated (*C, N, B*) and phosphorylated (*C*^*∗*^, *N* ^*∗*^, *B*^*∗*^). The transition between these two states depends on level of active ERK (*E*), which switches reversibly to it’s inactive form from the total constant pool of ERK *E*_max_. *k*_+_ and *k*_*−*_ are the rates of transition from inactive to active forms of ERK and vice-versa. Equation (1) describes the dynamics of ERK signals. *k*_+_ = *k*_endo_(*x*) + *k*_opto_(*x, t*), where *k*_endo_ has an endogenous profile from pole to the middle of the embryo and *k*_opto_ represents the optogenetic activation of ERK. In the model, ERK levels are same in the nucleus and cytoplasm. Equations (2)-(7) describe the dynamics among two states and three localization of Cic. The rate of phosphorylation of Cic is proportional to active ERK (*E*); *k*_p_ is the rate constant of phosphorylation, and *k*_d_ is the rate constant of dephoshorylation. We used the endogenous level in the pole of the WT embryo as the unit signaling level of ERK. Cic is synthesised with a volumetric rate of *q* and degrades in the cytoplasm, with the rate constant of *k*_deg_. The rate constants of degradation in cytoplasm are independent of the phosphorylation state of the molecule [14]. The rate constants of import and export are different for the unphosphorylated 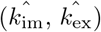 and phosphorylated 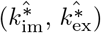 states. Similarly, affinity to DNA is also different for the unphosphorylated (*k*_b_) and phosphorylated 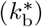 states. However, for the sake of simplicity we assumed the rate constants of unbinding to be same 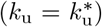 and rate of binding to DNA of unphosphorylated molecule to be significantly higher than the phosphorylated ones 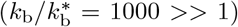. *S*^max^ is the total number of Cic binding sites in the *Drosophila* genome and 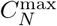 is the maximum concentration of nuclear Cic in nuclear cycle 14.

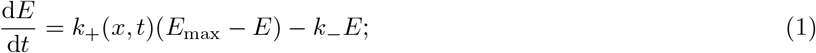

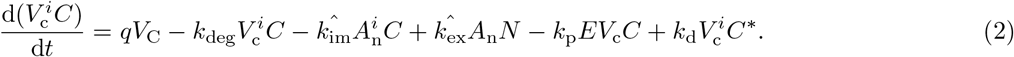

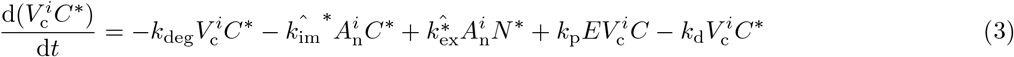

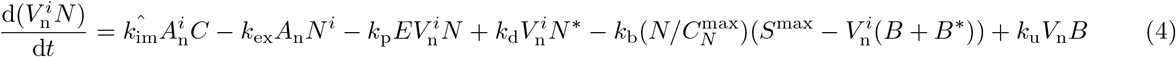

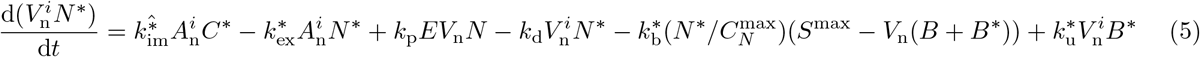

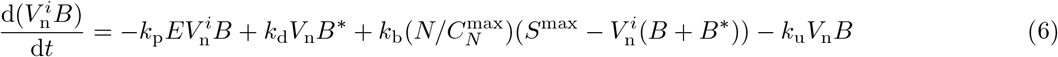

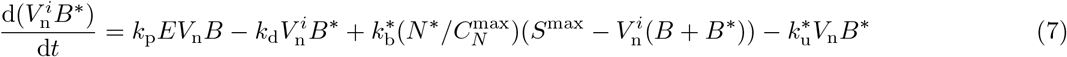

We non-dimensionalized all the concentrations by (*q/k*_deg_) and time as *τ* = *k*_deg_*t*. In cycle 11, we initialized the concentration of all the Cic pools as zero, whereas, in cycle 12-14 we distributed the total pool of unphosphorylated and phosphorylated Cic in the cytoplasm of two daughter cell, as the nuclear membrane breaks apart during mitosis. We used the geometric parameters 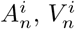 and 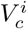 in nuclear cycle (*i*) 11-14 from [27]. For a given set of parameters, we calculated 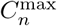 as the maximum value of *N* + *N* ^*∗*^ by setting 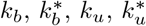 to zero and treating the bound and unbound pools together.

In the transcription, where Cic target flips between a transcriptionally active and inactive state (Fig. 1b). The rate constant of activation is *k*_on_, which is proportional to the probability of finding all the sites not bound by Cic, a non-linear function of fraction of Cic sites bound (*f*_b_), defined as 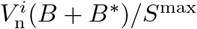 (Fig. 1b).

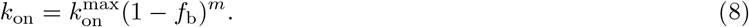

Deactivation happens with the rate constant *k*_off_. If transcription is off in a nucleus in time *t*, it switches to the on state in the next timestep at *t* + Δ*t*, with a probability exp(*−k*_on_(*t*)Δ*t*). Similarly, if transcription is on in a nucleus, it switches to the off state with a probability exp(*−k*_off_ Δ*t*).

### Parameter Estimation

The parameters for the tier 1 of the model were estimated by minimizing the difference of radial distribution function, a statistical mechanics descriptor that quantifies the structural order of a point pattern [20]. There are 12 free parameters in the rest of the model (Supplementary Table 1). In this 12 dimensional space, we started from an initial guess **p**_**0**_ chosen randomly in the pre-specified range (Supplementary Table 1). Then we ran the Markov Chain Monte Carlo (MCMC) simulations for *M* steps, where in each step *j* the following operations were done (Supplementary Code, [28]):

- The normalized error from each experiment ϵ_*kj*_ is calculated as 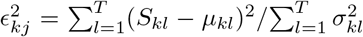, where, *µ*_*kl*_ and *σ*_*kl*_ are the mean and standard deviation of the *l*^th^ datapoint of the *k*^th^ experiment and *S*_*kl*_ is the equivalent model output.
- The total error of the simulation *E*_*j*_ is evaluated as the norm of a vector of normalized errors from all the experiments– 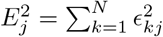.
- A vector **V** of length *δ* of the same dimension as the parameter space is chosen in a random direction.
- The parameter vector **p**^*g*^ is updated such that the *m*^th^ parameter is 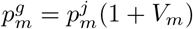.
- The total error *E*^*g*^ is calculated for the parameter vector **p**^*g*^.
- If *E*_*g*_ *< E*_*j*_, the movement in the parameter space is accepted with a probability 1, else, the movement is accepted with a probability 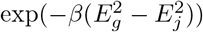.

As our initial guesses were uniformly dispersed in the parameter space, we chose *β* = 500 *>>* 1 and *δ* = 0.1 to find the local minimum around each initial guess. We present results from 1000 simulations, each of which ran for *M* = 2000 steps, when the error becomes steady (Supplementary Figure 7). In Fig. 2a-d, we plot the relative frequency of each pair of parameters in a bin in logarithmic scale.

### Comparing data and model predictions

In every step of the MCMC simulations, we solved equations (1)-(7) for 4 nuclei–a nucleus at the middle of a wild type embryo (*E* = 0), a nucleus at the pole of a wild type embryo (*E* = 1), a nucleus at the middle of the embryo with a step activation of ERK in cycle 14, and a nucleus at the middle of the embryo with a pulse activation of ERK of 5 minute duration in cycle 14 (*E* changes dynamically). We elaborate below how we compare the model output for these nuclei to the experimental data below.

#### Live Cic imaging

Though there are four pools of Cic in the nucleus, the live imaging of Cic-sfGFP doesn’t distinguish between the proteins bound and unbound to DNA or it’s state of phosphorylation. So, for comparison with the experimental data, we define the total dimensionless concentration of nuclear Cic as 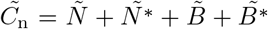, where 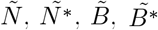 are dimensionless form of the respective concentrations. The nuclear Cic intensity obtained from live imaging are in arbitrary units. So, we further divide the non-dimensional nuclear Cic concentration 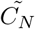 from the model by the concentration of cytoplasmic Capicua (*C* + *C*^*∗*^) at time *t*_0_, when the synthesis of Cic stops in cycle 14. We normalized the fluorescence intensity of Cic such that the maximum value of Cic intensity in cycle 14 in the middle of the embryo is 9, the experimentally reported ratio of nuclear and cytoplasmic Cic in cycle 14 [14]. In this way, we effectively normalized both the experimental and the computational time-series of nuclear Cic by the typical cytoplasmic concentration of Cic at cycle 14 (Fig. 1c). For comparing the ratio of nuclear Cic at the middle and pole regions, we found the ratio of 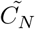 at time *t*_0_ in cycle 14 and compare it with the results from [17] (Supplementary Figure 2a).

#### DNA binding

The ChIP-Seq data reveals the genome-wide binding of Cic. Although this data is from the entire embryo, endogeneous ERK is active in less than 10 % of the embryo. So, we assumed the data to be reflective of nuclei without any endogeneous ERK signal such as one in the middle of the embryo within experimental error (≈ 20%). Furthermore, to compare with the model output *f*_*b*_, the fraction of Cic sites bound, we normalized all the experimental ChIP-Seq data to the wild type data, assuming all sites are bound in the middle of the wild type embryo in cycle 14.

#### Transcriptional response

The experimental data for transcriptional activity in the middle of the embryo in response to optogenetic activation is the fraction of nuclei transcribing in a given area [17], which is an ensemble property. However, during the MCMC optimization, we ran the simulations on a single nucleus on the middle of the embryo. To extract ensemble statistics out of this, we found the fraction of nuclei *f*_*t*_ transcribing in the middle of the embryo by solving the following equation,

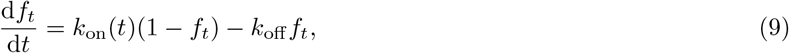

where, we calculated *k*_on_ as a function of *f*_*b*_ (equation 8) for the nucleus situated in the center of the nucleus. The initial condition is *f*_*t*_ = 0 after the mitosis. This comparison is based on the assumption that *E*, the only input to the model is constant in the imaging region, which is evident from the experimental ERK profiles [14, 16]. This predictions of this deterministic model are close to the predictions of a stochastic description of transcription (Supplementary Figure 8).

## Data and code availability

The code for the MCMC Simulation, data used to run the code and code for generating ensemble predictions are available in [28]. Source data for figure 1 and 2 are available with the manuscript.

## Acknowledgements

We thank Michael Frenklach, Eric Wieschaus, Trudi Schüpbach, and all members of the Shvartsman lab for helpful discussions. We thank Lucy Reading-Ikkanda for graphic design of the figures. We thank Gary Laevsky and Molecular Biology Core Confocal Microscopy Facility for imaging support. We thank Liu Yang for the ELISA experiment. We thank Nareg J.-V. Djabrayan and Gerardo Jimenez for the synthetic reporter CZC. We thank the Lewis Sigler Institute of Integrative Genmics for computational resources. The research was supported by the HD085870 grant from the NIH.

## Author Contributions

S.E.K. and A.L.P. carried out all the experiments. S.E.K., S.D. and A.L.P. analyzed the data. S.Y.S, S.D. and S.E.K. designed the model. S.D. implemented the model and performed the simulations. S.D., A.L.P., S.E.K., and S.Y.S. wrote the manuscript

