## Supplementary Figures 1-8 and Supplementary Table 1. for "From heterogeneous datasets to predictive models of embryonic development"

### From heterogeneous datasets to predictive models of embryonic development: Supplementary Material

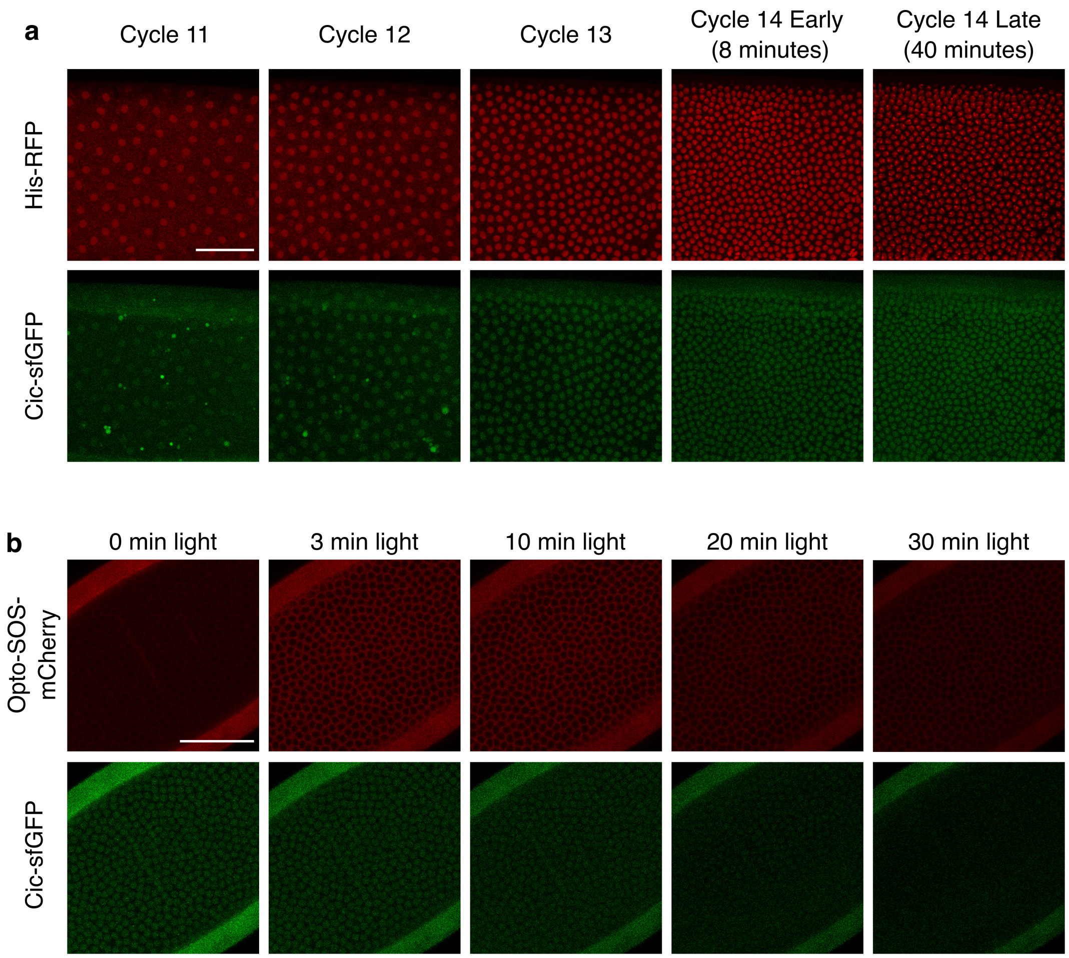


Figure 1: (a) Confocal images in the middle of the syncytial *Drosophila* embryo containing histone-RFP nuclear markers (red) and Cic-sfGFP (green) markers in a wild type embryo from cycle 11 to 14. (b) An embryo in early cycle 14 expressing cic-sfGFP (green) and the optogenetic ERK-activation tool optoSOS, which is tagged with the fluorescent protein mCherry (red). The time stamps indicate the duration of optogenetic stimulation. The scale bar denotes a length of 50 $\mu$m. Both the imaging in (a) and (b) were repeated $N = 3$ times each.

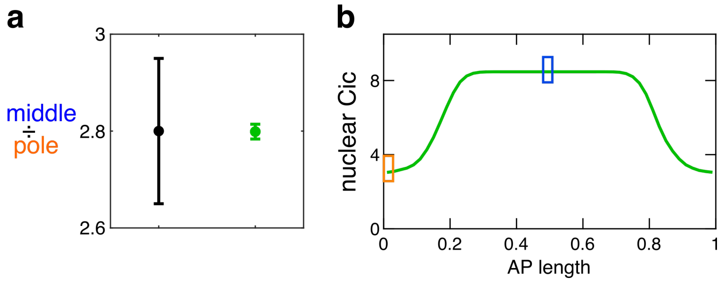


Figure 2: (a) Ratio of the Cic level at the middle versus the level at the anterior pole of the embryo. The mean and standard deviation from the experiment (black) and simulations with the parameter set from MCMC optimization (green) are shown. The black dot denotes experimental mean and the errorbar denotes the standard deviation across all experiments. (b) Simulated anterior to posterior (AP) profile of nuclear Cic level for one of the parameter sets given an ERK profile constructed so that the intensity is 1 at the middle and decays to zero at the poles. The blue and orange rectangles denote the locations of the points on the curve used to report the middle to pole Cic level ratio in (a).

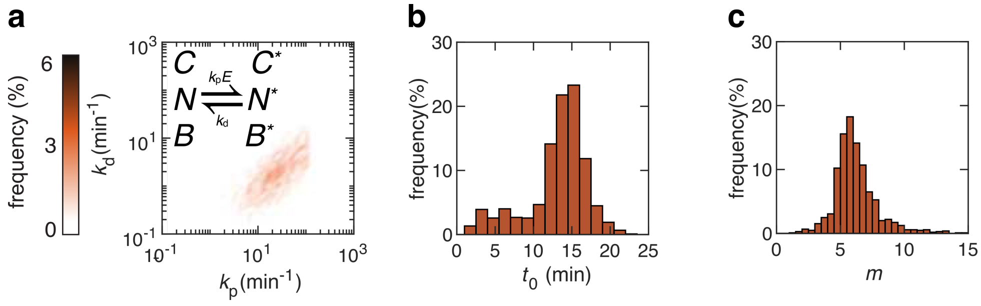


Figure 3: The distribution of the rate constants for phosphorylation ($k_p$) and dephosphorylation ($k_d$) (a), the time when synthesis of Cic stops in cycle 14 ($t_0$) (b), and the cooperativity parameter ($m$) (c) obtained from MCMC simulations.

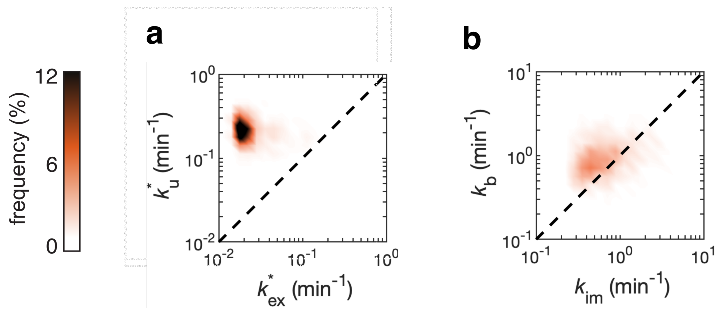


Figure 4: Two dimensional projections of parameters obtained from the MCMC simulation in $k^*_{\mathrm{ex}}$-$k^*_{\mathrm{u}}$ (a) [rate of unbinding vs. the rate of export of phosphorylated Cic] and $k_{\mathrm{im}}$-$k_{\mathrm{b}}$ (b) [rate of import vs. the rate of binding of unphosphorylated Cic] planes. The dotted lines represent the diagonals in these planes. We observe that the unbinding of phosphorylated Cic from DNA is significantly faster (above diagonal) than the export of phosphorylated Cic. The rate of binding of unphosphorylated Cic to DNA is also slightly faster (above diagonal) than the import of the same molecule in the nucleus. Both of these data suggest that nucleocytoplasmic shuttling is a slower process than the binding-unbinding of Cic to the genome.

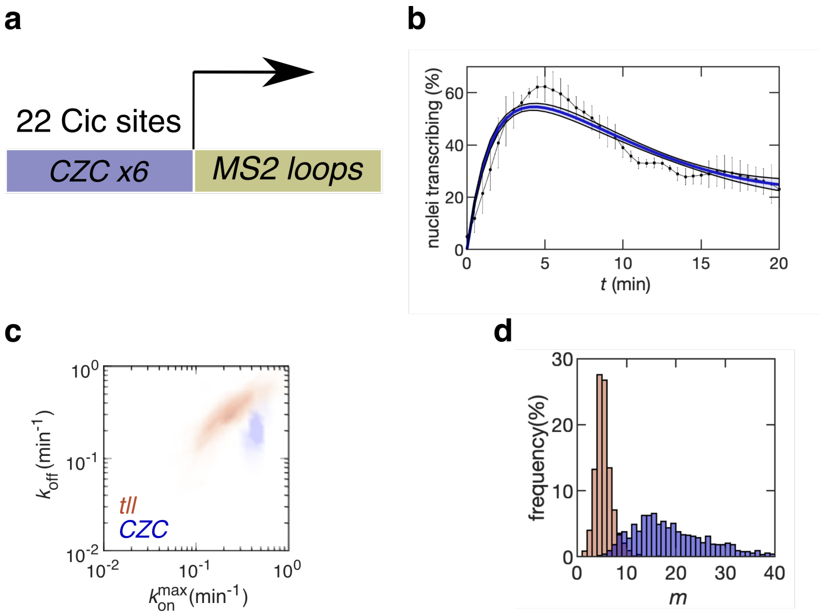


Figure 5: (a) Schematic of a synthetic reporter *CZC* containing 22 binding sites for Cic. (b) Transcriptional activity of *CZC* (black dots with errorbar), in the middle of an embryo stimulated with a direct optogenetic ERK signal (data from $N$=3 samples), and model prediction for the best fit parameters (blue). The black dots and the errorbars denote the experimental mean and standard deviation across all experiments. The light shaded region represents the standard deviation over all parameter sets. New parameters were learned using this experiment as an additional dataset, along with the other datasets mentioned in the main text. The parameters related to transcription of the synthetic reporter were kept free as they may be different from the reporter for *tll*. The rate of turning on ($k_{\mathrm{on}}^{\mathrm{max}}$) and turning off of transcription ($k_{\mathrm{off}}$) (c) and the cooperativity parameter $m$ (d) for *CZC* (blue) compared to *tll* (orange). The predicted $k_{\mathrm{on}}^{\mathrm{max}}$ for *CZC* is significantly higher than *tll*, suggesting the synthetic reporter exhibits stronger expression. The distribution of $m$ for *CZC* is shifted rightwards, which reflects that *CZC* has more Cic binding sites than *tll*. A more detailed model of transcription factor binding reflecting the 3D geometry of the genome may help to closely estimate the actual number of Cic sites. The other parameters of the model remained the same when re-optimized including the *CZC* dataset.

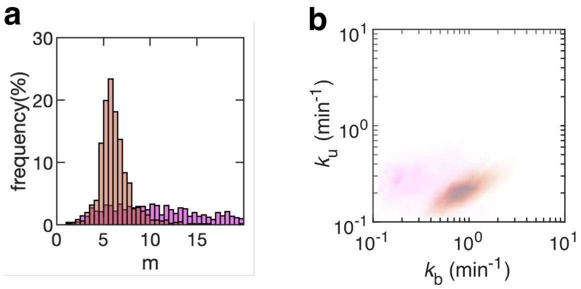


Figure 6: The distribution of rate constants of binding ($k_{\mathrm{b}}$) and unbinding ($k_{\mathrm{u}}$) to DNA (a) and the cooperativity parameter $m$ (b) if the ChIP data is excluded (purple) or included (orange). These parameters become constrained only when ChIP data was included, emphasizing the contribution of this particular experiment.

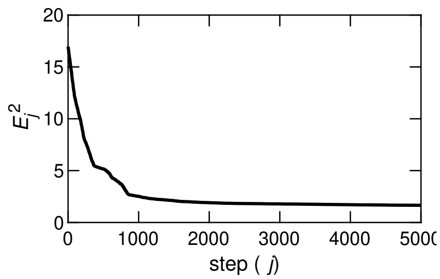


Figure 7: The square of the norm of the deviation of model outputs from the experimental results ($E_j^2$) as a function of Monte Carlo Markov Chain step ($j$). Here, we show that the error reaches a steady state within $j$=2000 steps. This is an average of 10 simulations.

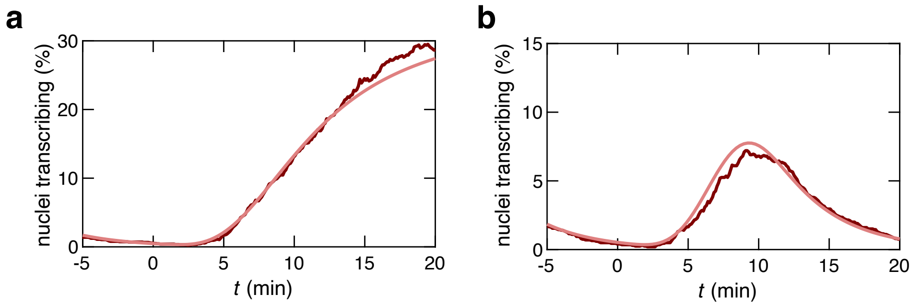


Figure 8: Comparison of the transcriptional response obtained from the solving equation (9) (light red line) and the transcriptional response obtained from stochastic simulations (dark red line) for a step (a) and pulse (b) signaling input in the middle of the embryo.

| | Fixed parameters | |
|---|---|---|
| parameter | physical significance | value |
| $q$ | rate of synthesis | eliminated by non-dimensionalization |
| $k_{\text{deg}}$ | rate of degradation | 0.09 $\text{min}^{-1}$ (ref. [14]) |
| $k_{\text{opto}}$ | rate of activation of light dependent ERK signal | 10 $\text{min}^{-1}$ (estimated from ref. [18])) |
| $k_{-}$ | rate of deactivation of ERK signal | 0.5 $\text{min}^{-1}$ (estimated from ref. [18]) |
| $E_{\text{ss}}(\text{opto})/E_{\text{ss}}(\text{pole})$ | ratio of steady state ERK signalling in an optogenetically activated embryo and at the pole | 2 (estimated from ref. [18]) |
| $V_n^i, A_n^i$ | area and volume of nucleus of in nuclear cycle $i$. $i = \{11, 12, 13, 14\}$. | data from (estimated from ref. [27]) |
| $\frac{S^{\text{max}}}{V_{\text{n}}^{14} C_{\text{n}}^{\text{max}}}$ | ratio of Cic site and maximum number of Cic molecule | 0.1 (data from ref. [17] and Liu Yang) |
| | Free parameters | |
| parameter | physical significance | range of initial guesses |
| $t_0$ | Time of stop of synthesis of Cic in cycle 14 | [5 15] min. (linear scale) |
| $k_{\text{im}} = \hat{k_{\text{im}}} A_n^{14}/V_n^{14}$ | rate of import of unphosphorylated Cic into nucleus in cycle 14 | [0.1 10] $\text{min}^{-1}$. (log scale) |
| $k_{\text{ex}} = \hat{k_{\text{ex}}} A_n^{14}/V_n^{14}$ | rate of export of unphosphorylated Cic from nucleus in cycle 14 | [0.1 10] $\text{min}^{-1}$. (log scale) |
| $k_{\text{im}}^* = \hat{k_{\text{im}}^*} A_n^{14}/V_n^{14}$ | rate of import of phosphorylated Cic into nucleus in cycle 14 | [0.01 1] $\text{min}^{-1}$. (log scale) |
| $k_{\text{ex}}^* = \hat{k_{\text{ex}}^*} A_n^{14}/V_n^{14}$ | rate of export of phosphorylated Cic from nucleus | [0.01 1] $\text{min}^{-1}$. (log scale) |
| $k_{\text{p}}$ | rate of phosphorylation in unit ERK signal | [1 100] $\text{min}^{-1}$. (log scale) |
| $k_{\text{d}}$ | rate of dephosphorylation | [1 100] $\text{min}^{-1}$. (log scale) |
| $k_{\text{b}}$ | rate of binding of unphosphorylated Cic | [0.01 1] $\text{min}^{-1}$. (log scale) |
| $k_{\text{u}}$ | rate of unbinding of unphosphorylated Cic | [0.01 1] $\text{min}^{-1}$. (log scale) |
| $k_{\text{on}}^{\text{max}}$ | rate of turning on transcription in absence of Cic | [0.01 1] $\text{min}^{-1}$. (log scale) |
| $k_{\text{off}}$ | rate of turning off transcription | [0.01 1] $\text{min}^{-1}$. (log scale) |
| $m$ | effective number of Cic sites near gene | [0 20] (linear scale) |

Table 1: The value of fixed parameters and and range of initial guesses of the free parameters of the model along with their physical significance.
